# VirB, a transcriptional activator of virulence in *Shigella flexneri*, uses CTP as a cofactor

**DOI:** 10.1101/2023.05.19.541425

**Authors:** Hammam Antar, Stephan Gruber

**Author notes:** Corresponding author, Stephan Gruber: S.G.

## Abstract

VirB is a transcriptional activator of virulence in the gram-negative bacterium *Shigella flexneri*. It is encoded by the large invasion plasmid, pINV, and is thought to counteract the transcriptional silencing mediated by the nucleoid structuring protein, H-NS. Mutations in *virB* lead to loss of virulence. Studies suggest that VirB binds to specific DNA sequences, remodels the H-NS nucleoprotein complexes, and changes DNA supercoiling. VirB belongs to the superfamily of ParB proteins which are involved in plasmid and chromosome partitioning often as part of a ParABS system. Like ParB, VirB forms discrete foci in *Shigella flexneri* cells harbouring pINV. Our results reveal that purified preparations of VirB specifically bind the ribonucleotide CTP. We show that VirB slowly but detectably hydrolyses CTP, which is mildly stimulated by the *virS* targeting sequences found on pINV. CTP and DNA binding promote VirB clamp closure. Curiously, DNA stimulation of clamp closure appears efficient even without *virS* sequences. These findings suggest that VirB acts as a CTP-dependent DNA clamp and may indicate that so far elusive factors might prevent offsite DNA clamping *in vivo*.

## Introduction

*Shigella flexneri* is the causative agent of the diarrheal disease shigellosis. It is a gram-negative bacterium that invades the epithelial lining of the intestinal tract [1]. *S. flexneri* contains a large ‘invasion’ plasmid pINV (∼220 kb) encoding several virulence factors including components of a Type III secretion system and its effectors [2]. pINV encodes a major virulence factor, VirB, an unconventional transcription regulator [3]. It is thought to counteract the transcriptional silencing of pINV mediated by the nucleoid associated protein, H-NS [4-8]. The molecular mechanisms underlying this de-repression are not entirely clear. Some putative models propose that VirB binds to specific DNA sequences and remodels the H-NS-DNA complexes making DNA more accessible for transcription (Fig. 1A) [9, 10]. A recent study moreover suggests that VirB triggers a change of DNA supercoiling of plasmid DNA *in vivo*, which is dependent on the VirB DNA binding sites, here designated as *virS* sites, thus proposing a new mechanism by which VirB could be offsetting the H-NS-dependent transcription silencing [11]. Another recent study showed that GFP-tagged VirB proteins forms discrete fluorescent foci in the cell that are dependent on the presence of the large invasion plasmid [12]. This focus formation is believed to stem from the accumulation of VirB on the target sites found on pINV, including one in the *icsP* promoter (herein called *virS*^*icsp*^). The *virS*^*icsp*^ site is sufficient for focus formation since a *Shigella* strain lacking pINV but carrying a plasmid with *virS*^*icsp*^ forms GFP-VirB foci. A crystal structure of the VirB middle domain (M domain) shows its helix-turn-helix motif bound to a DNA sequence found in the *icsB* promoter, another *virS* site (herein called *virS*^*icsB*^) [10]. However, other studies have suggested that VirB might bind different sequences, so the mechanism of targeting remains poorly understood [7, 8, 13].

**Figure 1:**
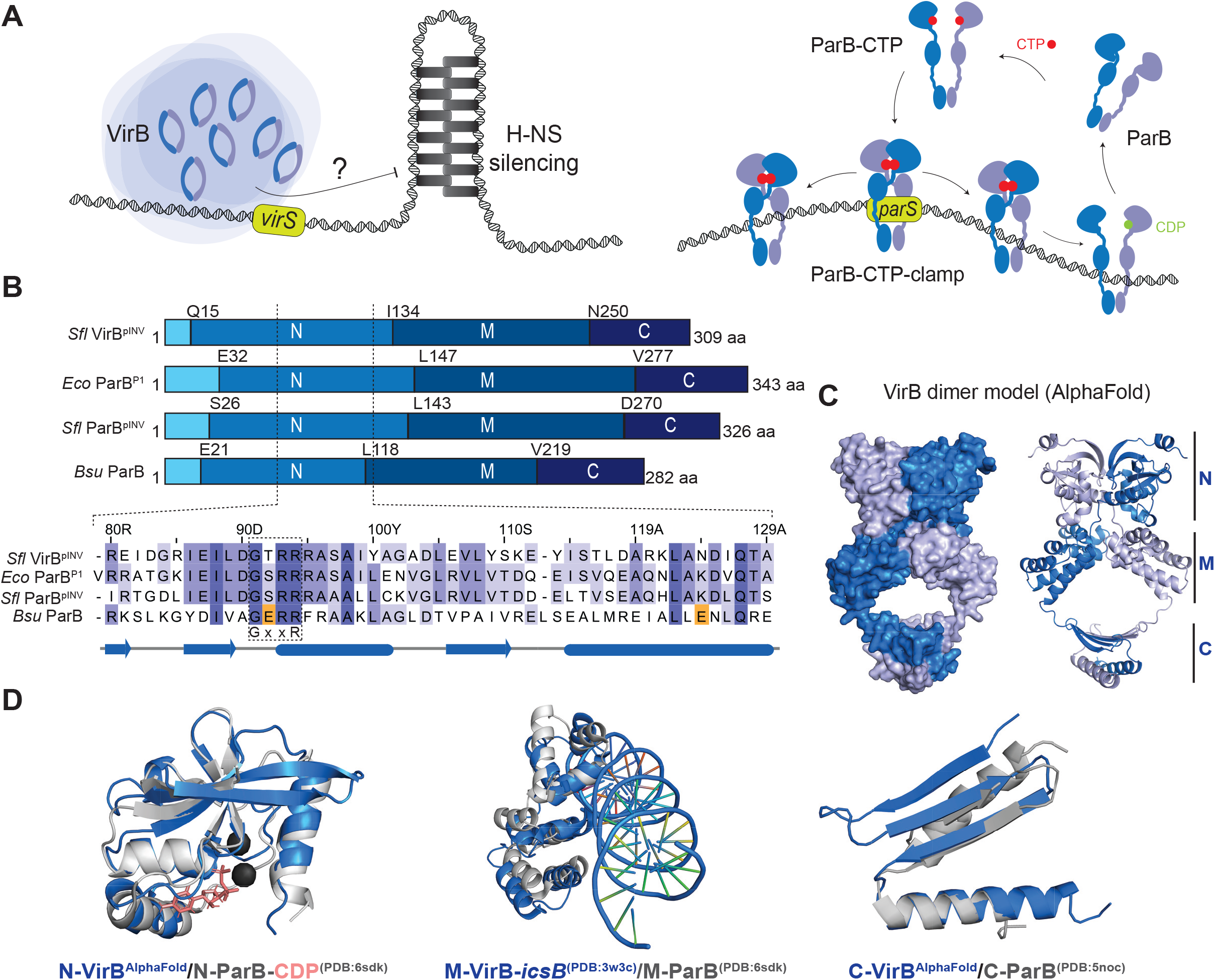
**(A)** the panel on the left shows a schematic representation of a model depicting the role of VirB in counteracting transcriptional silencing of pINV by H-NS. VirB is hypothesized to counteract the silencing effects of H-NS, a nucleoid associated protein. The exact mechanisms remain unclear, but potential models suggest that VirB may enhance DNA accessibility for transcription by sterically blocking H-NS or changing DNA conformation or supercoiling (not indicated). The panel on the right displays a schematic representation of ParB partition complex formation. In the absence of CTP binding, ParB exists in an open autoinhibited state. Upon binding to CTP and *parS* DNA, ParB self-inhibition is relieved resulting in the formation of ParB N domain dimer interface. This engagement of the N domain leads to the closure of ParB dimers, forming closed clamps that topologically entrap DNA and can slide onto *parS*-flanking DNA. Hydrolysis of CTP to CDP and inorganic phosphate destabilizes the N dimer engagement, promoting ParB release and turnover. This turnover mechanism prevents excessive spreading of ParB on DNA and allows for recycling of DNA-free ParB clamps. **(B)** Domain organization of *Sfl* VirB, *Ec* ParB^P1^, *Sfl* ParB^pINV^, and *Bsu* ParB (top panels). Sequence alignment of a part of the N domain of the four proteins including the conserved GxxR motif. The acidic residues responsible for CTP hydrolysis in *Bsu* ParB are highlighted in orange. **(C)** AlphaFold prediction of full-length *Sfl* VirB dimer displayed as surface and cartoon representation on the left and right panel, respectively. The prediction identifies three distinct domains (N, M, and C). **(D)** Separate superimposition of the three domains of *Sfl* VirB (blue) with the respective domain of *Bsu* ParB (gray). The N and C domains of VirB are AlphaFold predictions. The VirB M domain is from a crystal structure bound to the DNA targeting site, *virS*^*icsB*^ (PDB:3W3C) [10]. The three domains of *Bsu* ParB are from available structural data (PDB:6SDK for N- and M, and PDB:5NOC for C domain).

Based on sequence similarity, VirB belongs to the superfamily of ParB proteins which are normally involved in DNA partitioning as part of a ParABS system [7, 10, 14]. In fact, pINV encodes for two ParB homologues, which share 39 % sequence identity with one another, and 41 and 58 % identity with the *E. coli* P1 prophage ParB protein and somewhat lower similarity with chromosomally encoded ParB proteins (Fig. S1A). The gene encoding for the more closely related homologue of ParB^P1^ (58 % identity; herein referred to as ParB^pINV^) is located downstream of a ParA homologue likely promoting plasmid partitioning as a canonical plasmid ParABS system. The other one (41 % identity with ParB^P1^) is the transcription regulator VirB (Fig. 1B Fig. S1A).

ParABS systems comprise *parS* DNA sequences found near the origin of replication on the chromosome or plasmid, as well as the adaptor protein ParB and the partitioning ATPase ParA [15, 16]. ParA proteins form homodimers by binding ATP. ParA dimers associate with DNA in a sequence-unspecific manner.

Multiple ParB dimers load onto DNA at the *parS* sites, together forming a ParB/DNA partition complex (‘bacterial centromere’). This partition complex follows a ParA protein gradient on the bacterial chromosome to become equidistantly positioned within the cell. Once it interacts with ParA, it stimulates ParA ATP hydrolysis thus converting ParA dimers into monomers that dissociate from chromosomal DNA [17-19]. Following this diffusion ratchet mechanism, ParABS promotes chromosome partitioning and faithful DNA segregation [20].

In addition to being a DNA binding protein, ParB protein uses the unusual ribonucleotide cofactor CTP to mediate its functions in chromosome partitioning (Fig. 1B) [21-23]. ParB is an enzyme that binds and hydrolyses CTP using conserved motifs in the N-terminal domain (‘N domain’). Two CTP molecules are sandwiched between two N domains of a given ParB dimer [21-23]. *parS* DNA binding greatly stimulates the formation of a N domain dimer interface by relieving ParB self-inhibition [24]. This N-domain engagement turns ParB dimers into closed clamps that topologically entrap DNA and are thus able to slide onto the *parS-*flanking DNA covering up to 15 kilobases large regions around *parS*. On the other hand, hydrolysis of CTP to CDP and inorganic phosphate destabilizes N-domain engagement allowing for ParB turnover thereby preventing the excessive spreading on DNA and recycling any DNA-free ParB clamps [24-26]. Moreover, ParB clamps have been suggested to recruit other ParB dimers, in a CTP-dependent manner, to load on DNA distal from *parS* [27]. Altogether, CTP-dependent loading and one-dimensional sliding of ParB as well as dimer-dimer recruitment is thought to allow for focus formation near the origin of replication (partition complex) to support faithful chromosome segregation.

In this paper, and in light of recent discoveries, we aimed to unpack the similarities and differences between ParB and VirB proteins. We highlight the strong structural and biochemical resemblance between VirB and ParB. We show that VirB specifically binds and hydrolyses CTP. Using site specific BMOE-crosslinking, we show that CTP promotes VirB clamp closure *in vitro*, and that this closure is stimulated by specific and nonspecific DNA alike. It is thus conceivable that so far undiscovered factors promote site specific targeting of VirB (and maybe also ParB).

## Results

### AlphaFold prediction of VirB shows strong resemblance to ParB protein structure

AlphaFold predicts protein structures with reasonable confidence. AlphaFold predictions of VirB dimers display a closed, clamp-like configuration that resembles the organisation proposed for ParB dimers based on three separate ParB domains (N, M, and C domains) (Fig. 1C) [22, 23, 26, 28]. We superimposed the VirB model with the crystal structure of the *Bsu* ParB N domain bound to CDP (PDB: 6SDK). The superimposition showed close resemblance between the domains also revealing a pocket in VirB that may accommodate a ligand (Fig. 1D). This pocket is formed by sequences including the GxxR motif that is known to support CTP binding in ParB (Fig. 1B and S1A). Published structures of ParB and VirB M domains (PDB: 6SDK and 3W3C, respectively) as well as the C domains (PDB: 5NOC for ParB and an AlphaFold prediction for VirB) aligned well (Fig. 1D). We conclude that VirB shares multiple features with ParB proteins (including in the nucleotide-binding domain), implying that it could form partition complex-like nucleoprotein structures similar to ParB.

### VirB binds and hydrolyses CTP

We next addressed the question whether VirB binds any of the four ribonucleotides *in vitro*. Full-length VirB was recombinantly expressed and purified by N-terminal tagging with GFP. After removal of the GFP tag by proteolytic cleavage, full-length VirB was isolated and used in the following assays. Figure S1B displays the Size Exclusion Chromatography (SEC) profile of the purified protein indicating a single peak. The peak elution fraction was collected and used for the following experiments. The purity of the protein was confirmed by SDS-PAGE analysis, which showed a single band corresponding to the molecular weight of VirB. Based on measurements with isothermal titration calorimetry, VirB bound CTP with relatively high affinity (Kd ∼1 μM), while binding of the other ribonucleotides was not detected in this assay (Fig. 2A).

**Figure 2:**
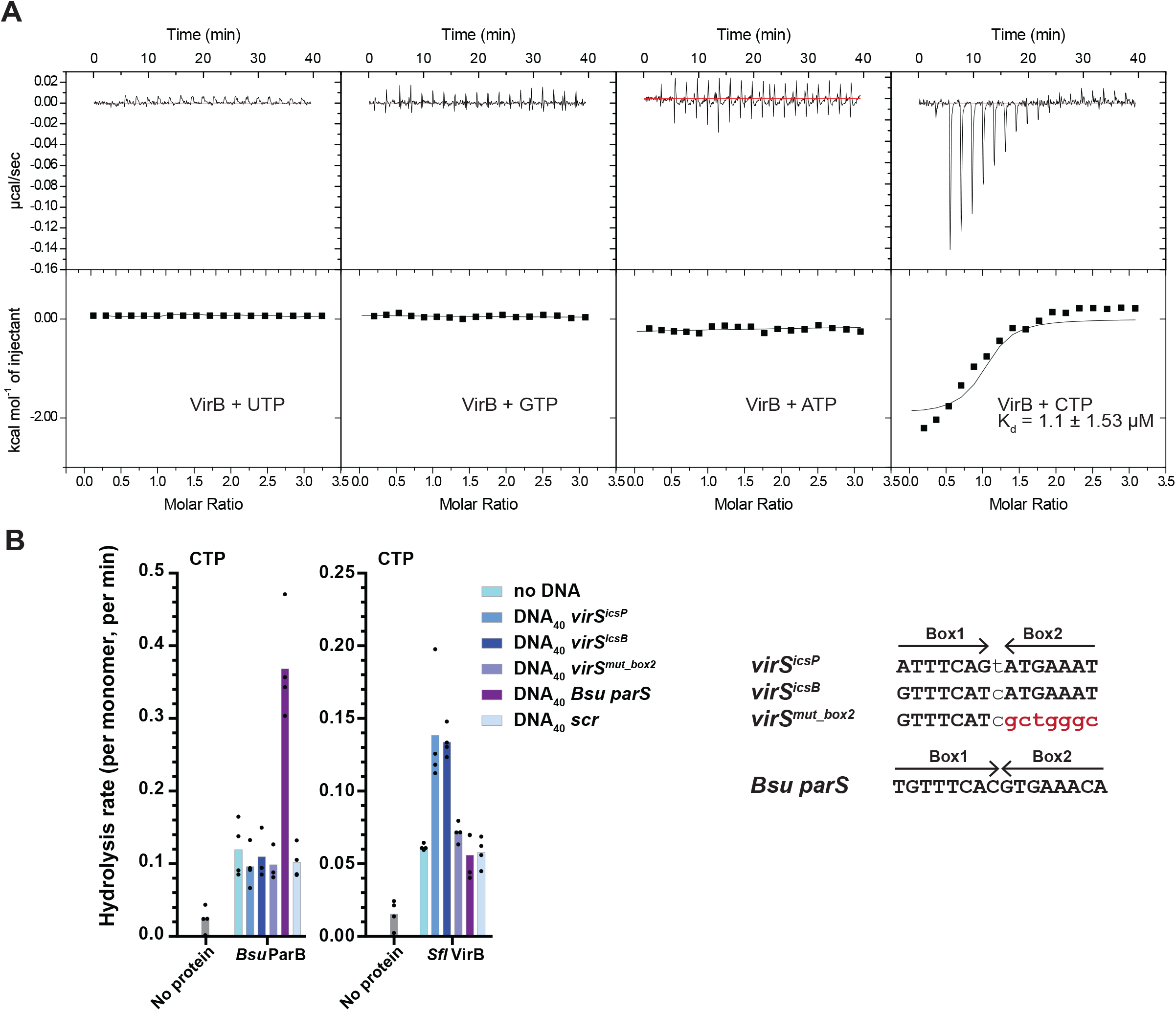
**(A)** VirB-ribonucleotide affinity measurements by isothermal titration calorimetry (ITC). A typical titration curve is shown. The K_d_ obtained from one experiment is provided. The interval indicates deviations of data points from the fit. **(B)** CTP hydrolysis rates by *Bsu* ParB (positive control) and *Sfl* VirB assayed by colorimetric detection of inorganic phosphate using Malachite Green Assay. Ten micromolar of protein was incubated with 1 mM CTP with or without 1 μM DNA_40_. Mean values calculated from four repeat measurements are plotted. Individual data points are shown as dots. The right panel indicates the sequence of the various DNA sites tested.

We then tested whether VirB can hydrolyse CTP (or one of the three other ribonucleotides) by measuring the release of free phosphate using Malachite green colorimetric detection. The protein showed low but noticeable basal levels of CTP hydrolysis in the absence of DNA (∼1 CTP hydrolysed/16 min per VirB monomer) (Fig. 2B). Upon addition of 40 bp DNA fragment containing *virS* sequences (*virS*^*icsp*^ or *virS*^*icsB*^), CTP hydrolysis was poorly but significantly stimulated (∼2-fold increase in hydrolysis rate: ∼1 CTP hydrolysed/7 min). The stimulation was not detected when mutated or scrambled *virS* DNA sequences were used (Fig. 2B). Usage of other ribonucleotide instead of CTP did not result in detectable levels of inorganic phosphate, suggesting that VirB specifically binds and hydrolyses CTP (Fig. S2A). Control reactions lacking protein showed that the low levels of inorganic phosphate detected with VirB are not contaminations (in the DNA and NTP preparations) (Fig. S2B). CTP hydrolysis is mildly stimulated by *virS* DNA sequences which we thus confirm to be specific recognition sequences for VirB protein.

### Efficient VirB clamp closure even in the absence of targeting DNA

We hypothesized that VirB proteins, like ParB, form DNA sliding clamps that self-load onto the specific target sequences *virS*. To detect the engagement of the N domains, *i*.*e*., closure of the VirB clamp, we employed site-specific cysteine crosslinking of purified VirB protein harbouring a cysteine mutation. Based on structural alignment with ParB dimers, we chose VirB residue Q15 to be mutated to cysteine for BMOE-crosslinking. Q15 falls at the axis of symmetry of the N domain dimer and should thus support robust cross-linking in the closed form of VirB (Fig. 3A). C5 was removed by mutagenesis to eliminate any unwanted cross-reactivity. Purified VirB(C5S, Q15C) protein exhibited comparable CTPase activity, indicating that the mutations did not significantly hamper protein folding or stability (Fig. S2A). In absence of ligands, a relatively small fraction of cross-linked VirB protein was detected (Fig. 3B) (∼18 %; lane 2). In the presence of CTP, more robust crosslinking was observed indicating that a significant fraction is found in a closed form (lane 3; ∼40 %) even in the absence of DNA. Addition of DNA further stimulated N domain engagement, particularly with *virS*^*icsp*^, *virS*^*icsB*^, and *virS*^*mut_box2*^ (lane 4-6; ∼70 %) and only slightly less so with other DNA sequences (lane 7 and 8; 55 %). Of note, significantly lower levels of closed VirB clamps were detected in the presence of DNA when CTP was lacking (lanes 9-13; ∼25-30 %). The results indicate that cofactors are not strictly required for VirB N-domain engagement (at least as measured by Q15C cross-linking), and that CTP alone can quite robustly support VirB clamp closure, unlike in canonical ParB proteins. Moreover, clamp closure is well stimulated by unspecific DNA (even at elevated salt concentrations (Fig. S2C)). Of note, other cysteine residues (endogenous C5 or engineered C5S, I30C) resulted in comparable outcomes albeit with overall reduced cross-linking efficiency likely owing to their larger Cys-Cys distances.

**Figure 3:**
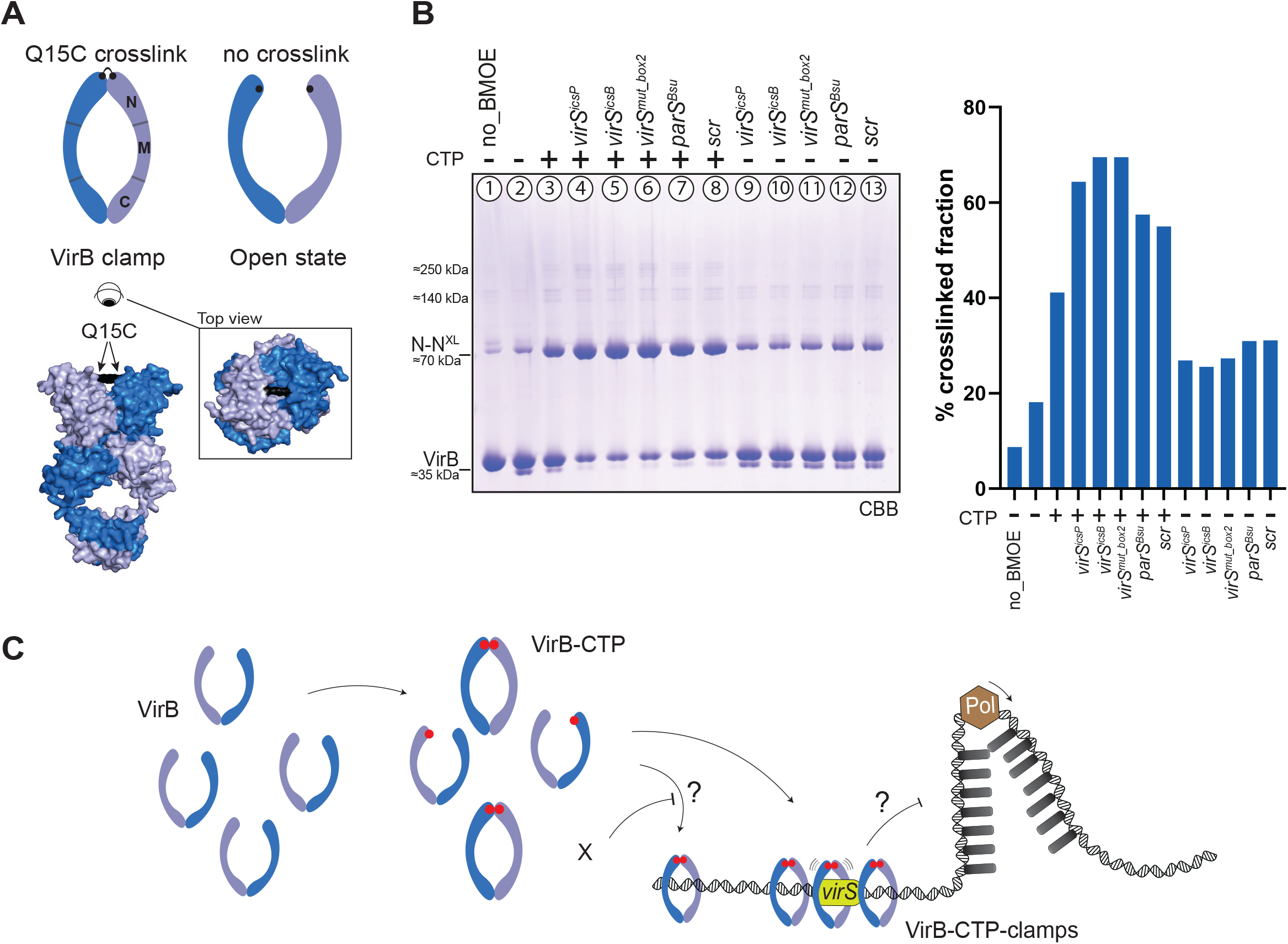
**(A)** Model of full-length VirB dimer with the mutated Q15C residue highlighted in black. Endogenous cysteine residue (C5) residue was replaced by a serine residue (C5S) to avoid unspecific crosslinking. The upper panel offers a visual representation of the crosslinked (clamp) and non-crosslinked (open) states of VirB. **(B)** Gel analysis of crosslinking products of purified VirB(C5S, Q15C). The different ligands tested are indicated. N-N^XL^ denotes crosslinked species of VirB. CBB, Coomassie Brilliant Blue. Quantification of cross-linked fractions is shown on the right panel. **(C)** Proposed model for VirB CTP binding and clamp closure. Upon CTP binding, VirB engages in the N domain and accumulate at *virS* sites and spread to neighbouring regions. X denotes a putative missing factor that could inhibit unspecific clamp closure and prevent VirB accumulation on unspecific DNA. VirB Focus formation at *virS* is hypothesized to counteract the silencing of virulence genes on pINV mediated by H-NS.

We conclude that CTP binding is required for robust VirB N-domain engagement and that it is stimulated by DNA in a largely sequence-non-specific manner, unlike in ParB gate closure, at least in our minimal reconstitution assay and under the reaction conditions used here. IF this observation directly translates to cell, it suggests that VirB clamps are formed away from *virS* sites and thus raises questions on how CTP binding may be linked to the accumulation of VirB specifically at the respective targeting sites on pINV.

## Discussion

In this paper, we show that VirB, a paralogue of ParB in *Shigella flexneri*, is a CTP-binding and -hydrolysing enzyme (Fig. 2). The usage of the unusual cofactor CTP is thus not restricted to ParB proteins and their roles in chromosome organization and partitioning, but also used by a transcription regulator, and potentially other cellular machineries. The nucleoid occlusion protein (Noc), a more closely related paralog of chromosomal ParB in firmicutes involved in the positioning of the cell division machinery in *Bacillus subtilis*, nucleates on the *parS*-like *nbs* sites using CTP [29]. It presumably does so by spreading on the DNA forming large nucleoproteins that hinder the assembly of the cell division machinery and direct it to the middle of the cell where division is supposed to occur.

VirB binds with a relatively high affinity (compared to ParB proteins) to CTP and hydrolyses it at a very low rate. The hydrolysis rate is mildly but specifically stimulated in presence of cognate DNA sites (*virS*^*icsp*^ and *virS*^*icsB*^) in our assay. Similar levels of stimulation have previously been observed with ParB proteins and *parS* sequences. In contrast, DNA stimulation of VirB N domain engagement is much less specific to *virS* DNA under the reaction conditions tested here. A significant fraction of the protein is found in a closed state even in the absence of DNA. This observation is puzzling because for ParB proteins, *parS-*stimulated clamp closure is believed to be a driving force for ParB accumulation near *parS* sites. In addition, ParB closure is thought to be the rate limiting step of CTP hydrolysis due to the formation of a self-inhibited state by ParB without *parS* DNA. Self-inhibition is relieved by *parS* DNA binding with the latter acting as a catalyst for clamp closure [24, 29]. This does not seem to be the case for VirB under the conditions tested here. VirB could be adopting an uninhibited state under our reaction conditions, making clamp closure efficient even in the absence of a catalyst, for example due to the lack of a putative clamp-closure inhibiting partner protein or cofactor. Consistent with this notion, we observed that VirB dimers entrap plasmid DNA *in vitro* regardless of the presence of *virS* sequences (Fig. S3).

**Table 1:**
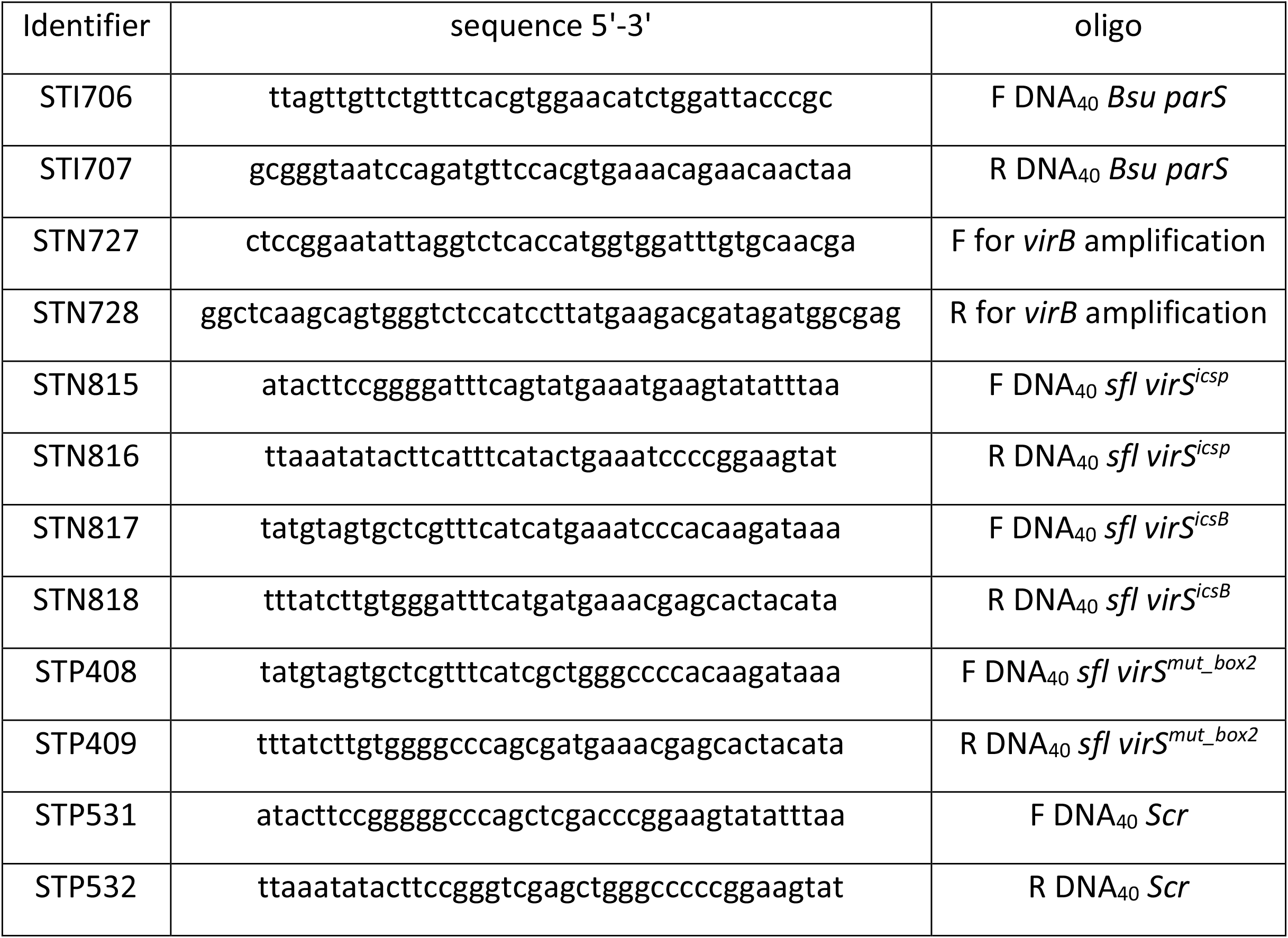
Primers used in this study

It is generally thought that CTP binding and hydrolysis by ParB protein is crucial for *parS* DNA loading and the formation of partition complexes, and in turn for the functions in chromosome partitioning. Our data suggest that CTP binding and hydrolysis also play a role in transcriptional regulation by VirB. The relevance and role of CTP binding and hydrolysis however remain to be determined. Constructing mutants of VirB that are defective in CTP binding or in CTP hydrolysis would be insightful in addressing the specific function of these steps of the CTP hydrolysis cycle. Curiously, the acidic residues contributing to CTP hydrolysis in chromosomal ParB proteins (GE_78_RRY/F and E_111_NLQR) are not found in VirB (or ParB^P1^ and ParB^INV^) (Fig. 1B and S1A). This implies that the mechanism of CTP catalysis is not fully conserved and possibly has been adopted during evolution.

Altogether, we propose that VirB uses CTP as a cofactor, potentially together with so far elusive accessory factors, to accumulate at high concentrations at the *virS* recognition sequences; the resulting VirB cluster is presumably needed to counteract the repressive function of H-NS proteins on the virulence plasmid for induction of virulence gene expression.

## Materials and Methods

### Expression and purification full-length proteins

**For *Bsu* ParB**, expression constructs were prepared in pET-28 derived plasmids by Golden-Gate cloning (as described [23]). Untagged recombinant ParB proteins were produced in *E. coli BL21-Gold (DE3)* grown in ZYM-5052 autoinduction media at 24 °C for 24 hours. **For *Sfl* VirB**, the protein was cloned from *Shigella flexneri* (DSM4782) obtained from DSMZ (German Collection of Microorganisms and Cell Cultures). The cloning process involved the use of specific primers designed to amplify the *virB* gene (See table of primers). Expression constructs were then assembled into pLIBT7 derived plasmids by Golden-gate cloning with an N-terminal GFP-tag [30]. GFP-tagged recombinant VirB proteins were produced in *E. coli BL21-Gold (DE3)* grown in TB-medium at 37°C to an OD (600nm) of 1.0 and the culture temperature was reduced to 24°C. Expression was initiated with the addition of IPTG to a final concentration of 0.4 mM and was allowed to continue overnight, typically for 16 hours. Purification of proteins was done as described before in [23]. In brief, cells were lysed by sonication in buffer A (1 mM EDTA pH 8, 500 mM NaCl, 50 mM Tris-HCl pH 7.5, 5 mM β-mercaptoethanol, 5 % (v/v) glycerol, and protease inhibitor cocktail (PIC, Sigma)). The first step **for ParB purification** involves adding Ammonium sulfate to the supernatant until it reaches 40% saturation and allowing it to stir at 4 °C for 30 minutes. The sample is then centrifuged, and the supernatant is collected. Additional Ammonium sulfate is added to the sample to reach 50% saturation, and it is allowed to stir at 4 °C for another 30 minutes. The pellet is collected by centrifugation and dissolved in buffer B (50 mM Tris-HCl pH 7.5, 1 mM EDTA pH 8 and 2 mM β-mercaptoethanol). **For VirB**, the first step involved running the supernatant on a homemade GFP column, and then the bound protein was proteolytically cleaved with an HRV-3C Protease overnight. **Both proteins** were subjected to the same purification protocol thereafter. The sample was additionally diluted with buffer B to a conductivity of 18 mS/cm and loaded onto a Heparin column (GE healthcare). The protein was eluted with a linear gradient of buffer B containing 1 M NaCl. Peak fractions were collected and diluted with buffer B to a conductivity of 18 mS/cm and loaded onto HiTrap SP columns (GE healthcare). A linear gradient of buffer B containing 1 M NaCl was used for elution. Peak fractions were collected and directly loaded onto a Superdex 200 16/600 pg column (GE healthcare) preequilibrated in 300 mM NaCl and 50 mM Tris-HCl pH 7.5. For cysteine mutants, 1 mM TCEP was added to the gel-filtration buffer.

### Isothermal titration calorimetry (ITC)

ITC measurements were done using MicroCal iTC200 (GE Healthcare Life Sciences). The device was cooled to 4 °C before use. All measurements were performed in a buffer containing 150 mM NaCl, 50 mM Tris/HCl (pH 7.5), and 5 mM MgCl2. Purified protein peak fractions from the Superdex 200 16/600 pg column were collected, diluted 1:1 with buffer containing 50 mM Tris-HCl pH 7.5 and 10 mM MgCl2 to bring the final buffer to 150 mM NaCl, 50 mM Tris-HCl pH 7.5 and 5 mM MgCl2 and directly used for measurements. Both the measurement cell and the injection syringe were thoroughly cleaned with the buffer. The measurement cell was filled with 280 μL of protein solution at a 30 μM monomer concentration, while the injection syringe was filled with buffer containing 0.5 mM NTP concentration or just buffer. The measurements were initiated after a 180-second delay, and the instrument settings were set to: reference power of 5 μcal/sec, stirring velocity of 1000 rpm, and “high feedback” mode. The raw data, expressed in kcal/mol, were presented as a Wiseman plot, and regression curves were calculated using a 1:1 nucleotide-to-protein monomer binding model when applicable. Origin software (GE Healthcare) was employed for fitting the measurement results using the equation:

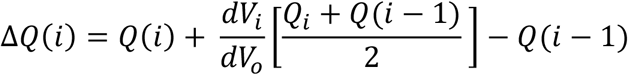

In this equation, *V*_*i*_ is the injection volume of ligand (nucleotides), *Vo* denotes the cell volume, *Q(i)* signifies the heat released from the *i*^*th*^ injection which is in turn calculated using the following equation:

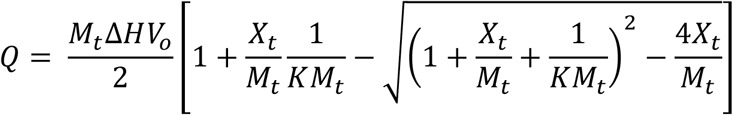

Here, *K* is the binding constant, *ΔH* is the molar heat of ligand binding, *Xt* refers bulk concentration of nucleotide, and *Mt* is the bulk concentration of ParB (moles/liter) in *Vo. K* and *ΔH* were estimated by Origin, and *ΔQ(i)* for each injection was calculated and compared to the measured heat. To refine the estimates of *K* and *ΔH*, standard Marquardt methods were applied, and iterative adjustments were made until no further improvement in the fit could be achieved.

### Measurement of NTP hydrolysis by Malachite Green colorimetric detection

NTP hydrolysis was measured as described in [24]. In brief. Mixtures of NTP (2x) with or without DNA_40_ (2x) and mixture of protein solutions (2x) were prepared in reaction buffer (150 mM NaCl, 50 mM Tris pH 7.5, 5 mM MgCl_2_) on ice. Equal volumes of each solution were mixed together (protein:ligand 1:1) using BenchSmart 96 (Rainin) dispenser robot and mixed through pipetting. Post-mixing, samples (containing 1 mM NTP, 1 μM DNA_40_, and 10 μM protein) were then incubated at 25 °C for 1 hour, and phosphate blanks were prepared in parallel. After incubation, samples were diluted four-fold by adding 60 μL of water, followed by mixing with 20 μL of working reagent (Sigma). The samples were then transferred to a flat-bottom 96-well plate. The plate was left to incubate for 30 minutes at 25 °C, after which the absorbance was measured at a wavelength of 620 nm. Absorbance values from the phosphate standard samples were used to plot an OD_620_ versus phosphate concentration standard curve. Raw values were converted to rate values using the standard curve, and absolute rates were determined by normalizing for protein concentration. Mean values and standard deviation were calculated from four replicates and presented as graphs on GraphPad Prism software.

### Preparation of 40-bp double stranded DNA

To generate 40-bp double-stranded DNA, two complementary oligonucleotide strands at a concentration of 100 μM each were combined in a 1:1 ratio. The resulting mixture was heated to 95 °C for 10 minutes and subsequently allowed to cool down to 25 °C.

### *In vitro* cross-linking

A 2x CTP solution was prepared with or without DNA_40_ in reaction buffer (composed of 150 mM NaCl, 50 mM Tris-HCl pH 7.5, 5 mM MgCl2), and these mixtures were allowed to sit at room temperature for 5 minutes. A 2x protein solution (in the same buffer) was added to the mixture to obtain the following final concentrations: 10 μM protein, 1 mM CTP and 1 μM DNA_40_. The samples were then incubated for an additional 5 minutes at room temperature before adding 1 mM BMOE. After another 5 minutes at room temperature, the samples were quenched with β-mercaptoethanol (23 mM final). Loading dye was added and the samples were incubated at 70°C for 5 minutes. Subsequently, they were loaded onto Bis-Tris 4-12 % gradient gels (ThermoFisher). The bands were stained with Coomassie Brilliant Blue (CBB), and the relative band intensity was quantified by scanning and semi-automated analysis in ImageQuant (GE Healthcare).

### Plasmid DNA entrapment assay

The plasmid entrapment assay was performed as described in [23] and similarly to *in vitro* crosslinking. The final concentrations of CTP, plasmid DNA or DNA_40_, and VirB protein were 250 μM, 50 nM, and 2 μM, respectively. Reaction mixtures were incubated at room temperature for 5 minutes and then treated with 1 mM BMOE for 5 minutes. After quenching with β-mercaptoethanol (23 mM final), the samples were divided into two halves. The first half was treated with 1x SDS loading dye for protein visualisation and incubated at 70°C for 5 minutes and loaded onto a WedgeWell Tris Glycine 4-12% gradient gel (ThermoFisher). Coomassie Brilliant Blue (CBB) staining was performed to detect proteins. The other half was mixed with 1x DNA loading dye and DNA detection involved running the samples onto 1% (w/v) TAE agarose gel containing ethidium bromide (0.5 μg/mL) and electrophoresed at 4°C, 10 V/cm for 1-2 hours. The agarose gel was then visualized using a Gel Doc XR+ (BioRad).

## Acknowledgements

We are grateful to members of the Gruber lab for stimulating discussions and comments on the manuscript and to Martin Thanbichler for sharing unpublished results.

---

**Figure S1:**
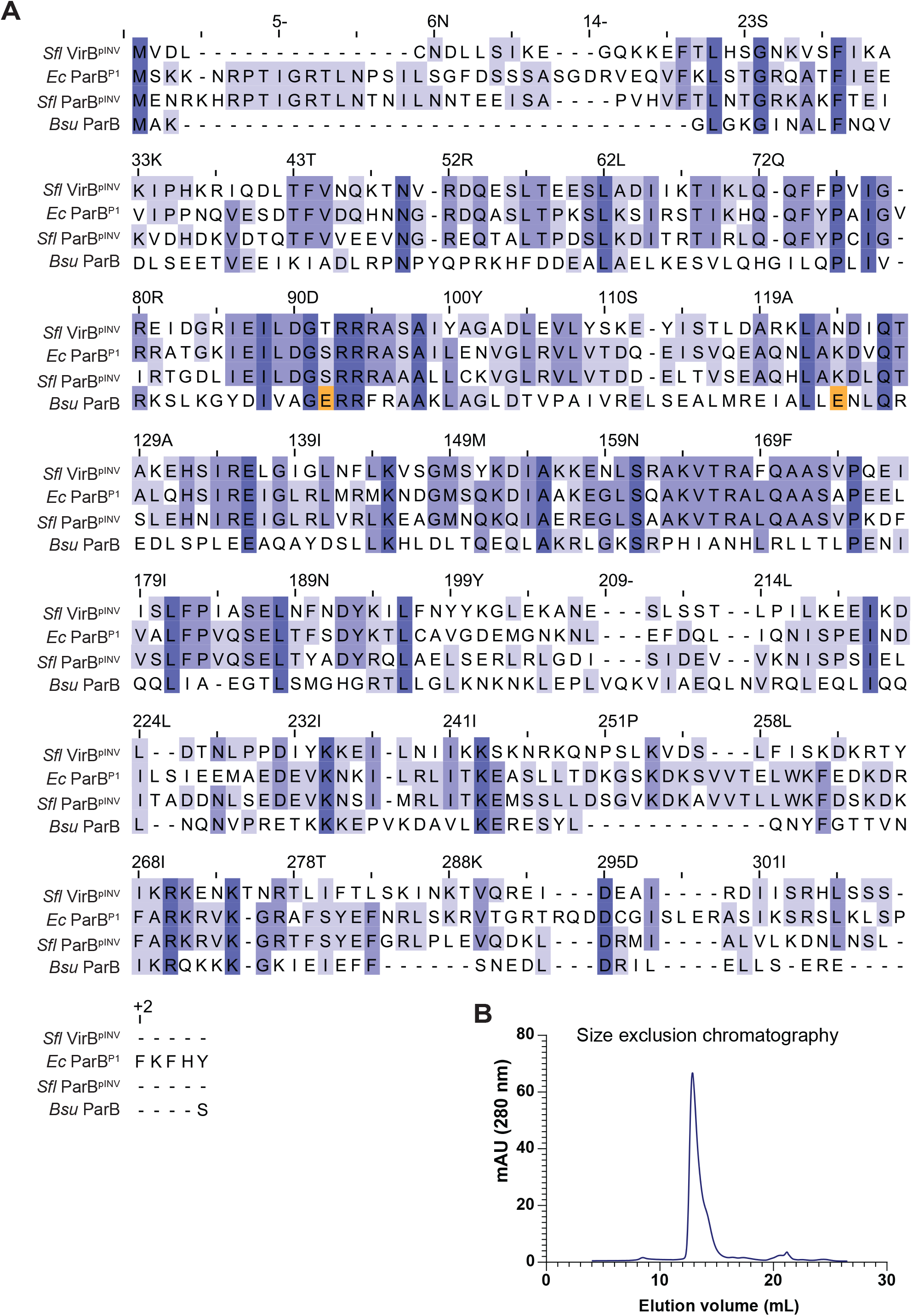
**(A)** Sequence alignment of full-length *Sfl* VirB^pINV^, *Ec* ParB^P1^, *Sfl* ParB^pINV^, and *Bsu* ParB. The sequence alignment was done on JalView and the residues are colour coded based on percentage identity. **(B)** Size-exclusion chromatography (SEC) profile of purified VirB protein. The SEC profile displays a single peak, indicating a homogenous protein sample.

**Figure S2:**
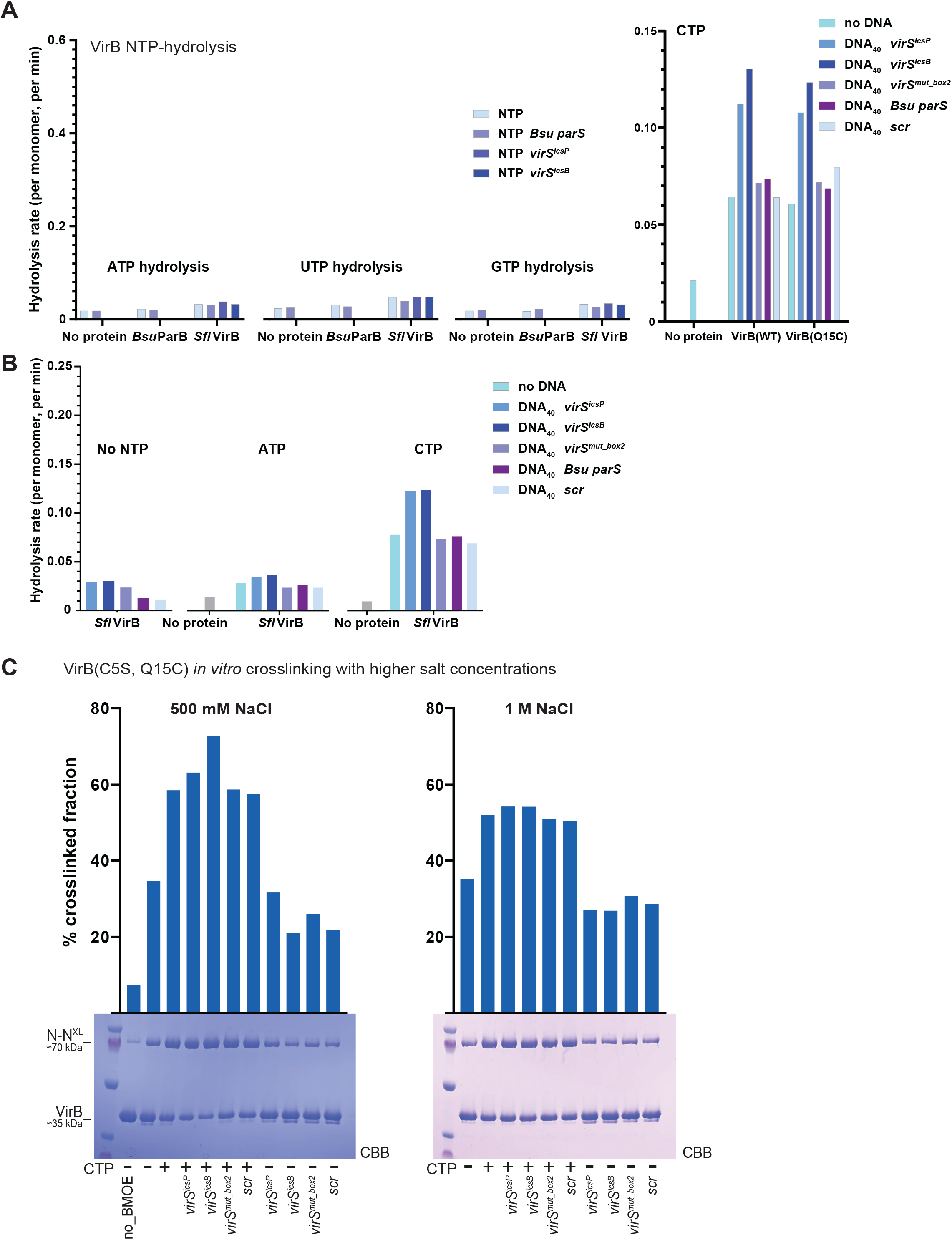
**(A)** Nucleotide hydrolysis by *Bsu* ParB and *Sfl* VirB(WT) and VirB(C5S, Q15C) measured by malachite green detection of inorganic phosphate. Data for CTP is also shown in Fig. 2B. Additional information can be found in the Materials and Methods section. **(B)** Detection of free inorganic phosphate via Malachite green assay. Same as in Fig. S2A. **(C)** Same as in Fig. 3B: Gel analysis of crosslinking products of purified VirB(C5S, Q15C) in higher salt conditions (500 mM and 1 M NaCl).

**Figure S3:**
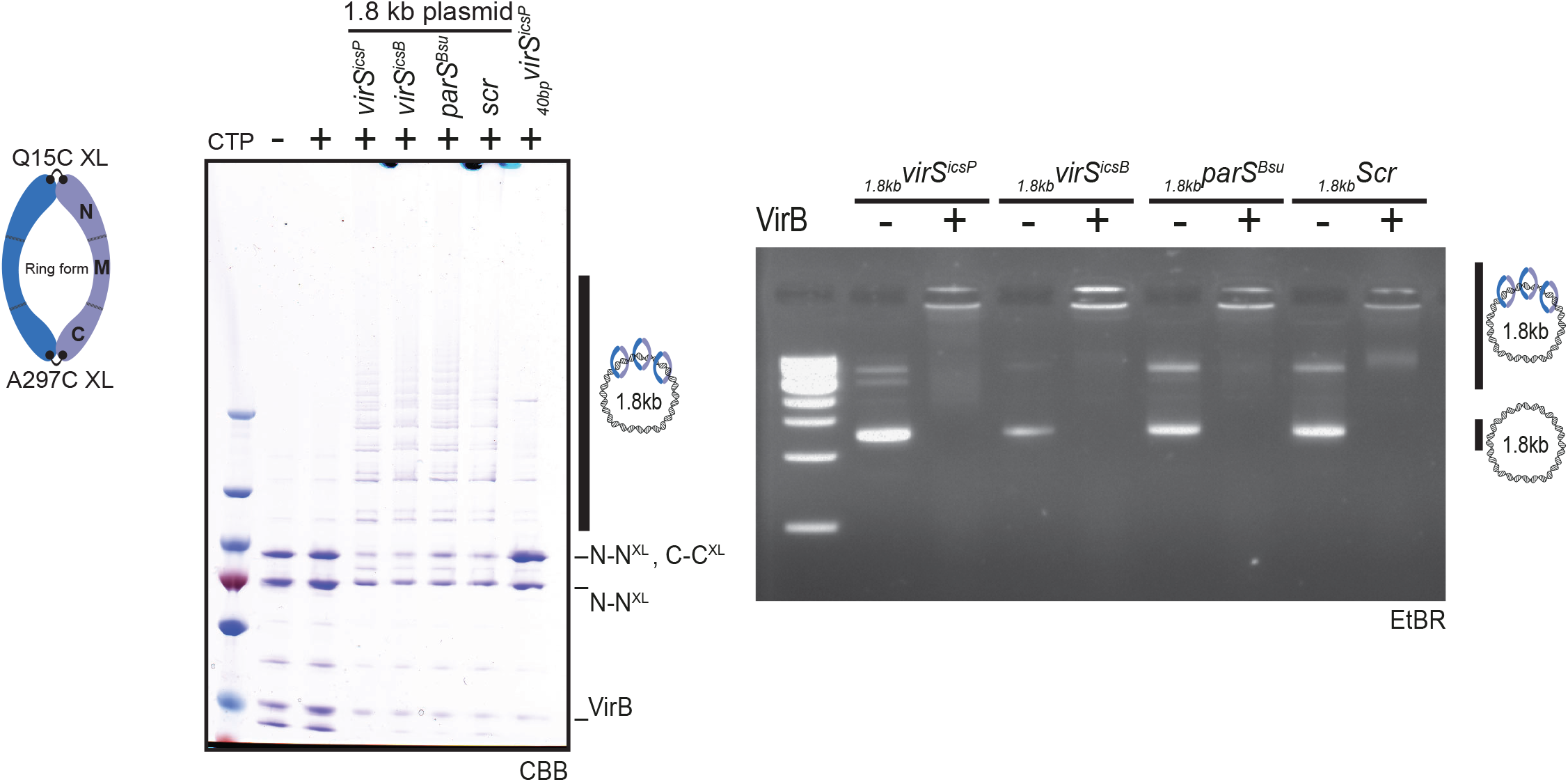
Plasmid entrapment assay by VirB(C5S, Q15C, A297C): polyacrylamide gel electrophoresis analysis of VirB protein species (CBB) and DNA species (ethidium bromide EtBr) from BMOE-cross-linked DNA loading reactions. The figure shows the entrapment of circular DNA by VirB, as observed in all four tested plasmids in the presence of CTP. A putative topologically entrapped circular plasmid by double crosslinked VirB dimers (VirB-XX) is marked.

